# A Bayesian Hierarchical Model of Trial-To-Trial Fluctuations in Decision Criterion

**DOI:** 10.1101/2024.07.30.605869

**Authors:** Robin Vloeberghs, Anne E. Urai, Kobe Desender, Scott W. Linderman

## Abstract

Classical decision models assume that the parameters giving rise to choice behavior are stable, yet emerging research suggests these parameters may fluctuate over time. Such fluctuations, observed in neural activity and behavioral strategies, have significant implications for understanding decision-making processes. However, empirical studies on fluctuating human decision-making strategies have been limited due to the extensive data requirements for estimating these fluctuations. Here, we introduce hMFC (Hierarchical Model for Fluctuations in Criterion), a Bayesian framework designed to estimate slow fluctuations in the decision criterion from limited data. We first showcase the importance of considering fluctuations in decision criterion: incorrectly assuming a stable criterion gives rise to apparent history effects and underestimates perceptual sensitivity. We then present a hierarchical estimation procedure capable of reliably recovering the underlying state of the fluctuating decision criterion with as few as 500 trials per participant, offering a robust tool for researchers with typical human datasets. Critically, hMFC does not only accurately recover the state of the underlying decision criterion, it also effectively deals with the confounds caused by criterion fluctuations. Lastly, we provide code and a comprehensive demo at www.github.com/robinvloeberghs/hMFC to enable widespread application of hMFC in decision-making research.

## Introduction

Every day we make a multitude of decisions: deciding when to cross the street, whether to add a newly discovered song to your playlist, or choosing between coffee or tea. Much effort has gone toward understanding the computational mechanisms underlying perceptual (1), value-based (2), risky (3), and social decisions (4). A common theme in these frameworks is that decisions are formed by comparing evidence to a decision criterion. Classical models of decision-making assume that observers generate an internal representation of the relevant information, typically referred to as “evidence” or the “decision variable” (5). To generate a binary choice, this decision variable is compared to an internal decision criterion. Due to the inherent noisiness of the brain or changes in internal states (e.g., changes in attention), repeated presentation of the same stimulus will not always lead to the same internal representation of that stimulus. Instead, multiple trials of the same stimulus produce a distribution of internal decision variables (fig. 1A). This inherent variability explains why participants make variable responses when repeatedly presented with the same stimulus.

**Figure 1:**
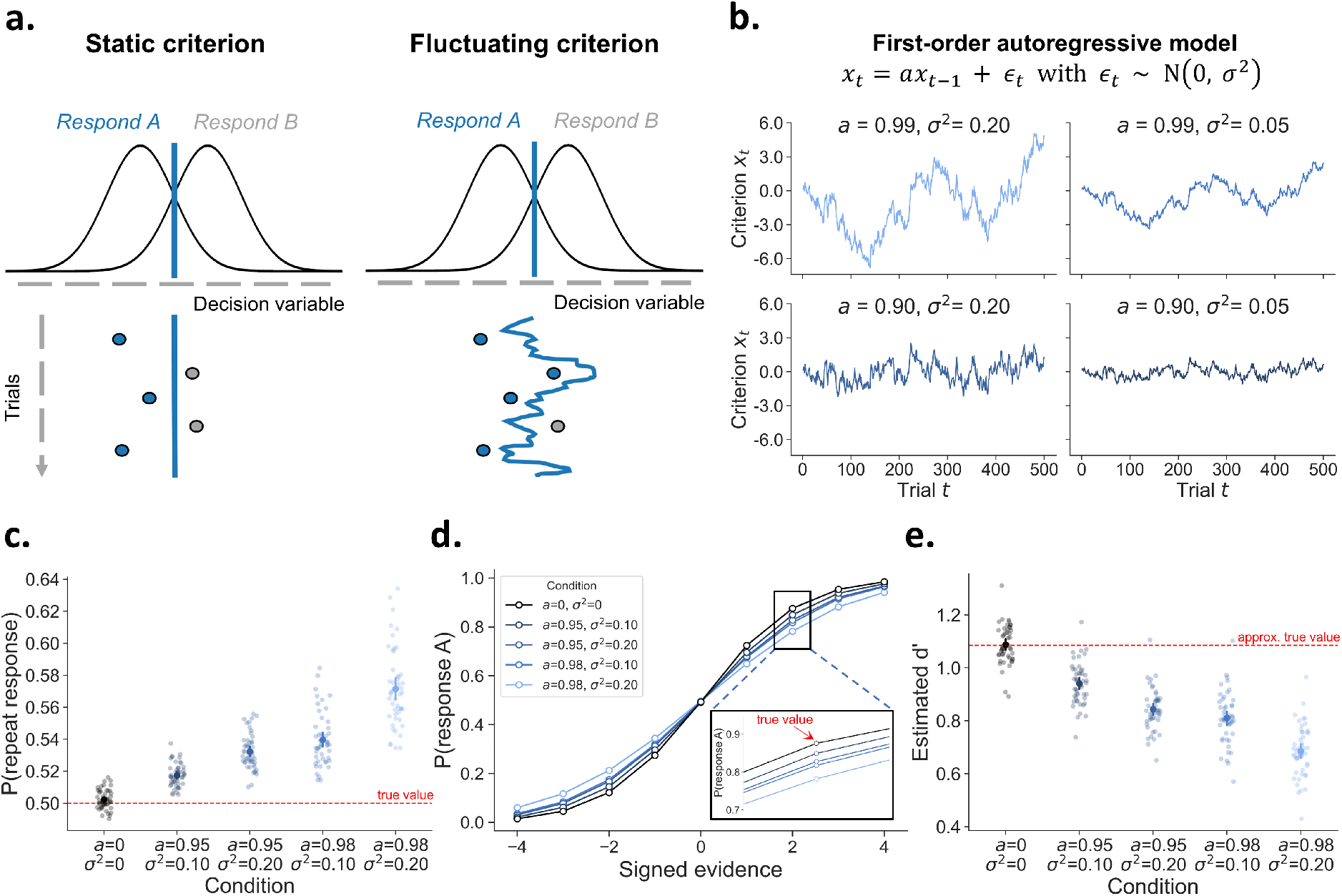
The consequences of fluctuations in decision criterion. **A)** Many theories of decision-making assume that choices are formed by comparing a decision variable to a criterion. According to signal detection theory, an observer will respond A or B, depending on whether the decision variable falls left or right to this criterion, respectively. Whereas this criterion is typically assumed to be fixed across a series of trials (static criterion), here we investigate the consequences of this criterion slowly fluctuating across time (fluctuating criterion). **B)** Fluctuations in criterion can be modelled as a time series using a first-order autoregressive model AR(1). An AR(1) model has two free parameters that control the temporal dependency (*a*) and the scale of the fluctuations (*σ*^2^). The four panels show how the criterion changes over time under different values of these two parameters. **C)** Simulations show that it is of critical importance to consider fluctuations in decision criterion. Despite the absence of an effect of previous response (true *β*_previous response_ = 0), they give rise to artificial choice history effects. **D)** Likewise, while the generative psychometric slope (true *β*_stimulus_ = 1.25) is constant for all conditions, fluctuations in decision criterion lead to an underestimation of the slope of the psychometric function. **E)** Fluctuations in criterion lead to an underestimation of *d*′, a popular signal-detection measure of sensitivity.

A key component in these models is the decision criterion, which allows to explain biases in decision-making. When observers choose one answer more often than another, their behavior can be modelled as the result of a shift in decision criterion, independent of their sensitivity to decision-relevant information (5). Empirically, the decision criterion depends on internal and external factors: for instance, it shifts in response to external feedback (6), expectancy (7), task instructions (8), monetary rewards (9), and the base rate of stimuli (10).

Although it is widely recognized that the decision criterion can be flexibly shifted depending on the environment, a common implicit assumption is that in the absence of experimental manipulations, this (potentially biased) criterion remains constant across a series of experimental trials. Although this assumption allows computational models to be simple and tractable, it may be overly simplistic. Indeed, the idea of trial-by-trial variability in the criterion can be traced back to early work (11; 12; 13), and has been noted often in the literature (14; 15). However, this line of work focuses mostly on the criterion distribution (mean and variance), rather than attempting to estimate the trial-to-trial trajectory of the fluctuating decision criterion.

More recently, the dynamical nature of cognitive processes has received renewed interest in cognitive and computational neuroscience (16). One reason for this trend is the availability of new computational tools to quantify trial-by-trial fluctuations from behavioral and/or neural data. Indeed, increasing evidence suggests that several computational parameters that are thought to underlie decision-making are not stationary, but rather fluctuate over trials (17; 18; 19; 20; 21). For example, Ashwood and colleagues (17) demonstrated that decision behavior in mice can be described by three discrete states persisting for 10 to 100 trials, each with different psychometric parameters. Other work has focused on modeling continuous trial-by-trial changes in the overall bias (i.e., intercept; (19)) and other parameters of a logistic regression model (21). Combining behavior with neural recordings, Cowley et al. (18) found that during a multi-hour session, monkeys’ hit and false alarm rate slowly drifted (i.e. suggesting a slow drift in the decision criterion), and co-varied with neural activity in area V4 and prefrontal cortex. In other work, Mochol and colleagues (20) showed that decision behavior in monkeys is influenced by fast (previous trial’s response and reward) and slow (moving window of 130 trials) biases, both encoded in the PFC, that are integrated into the current decision.

Despite much progress in this line of research, an important shortcoming in the current literature is that the few methods that are available to estimate the decision criterion at the *single-trial* level require too many trials and fix crucial parameters instead of estimating them. In this work, we propose a new model that solves these issues. First, we start with investigating the consequences of fluctuations in decision criterion. We do so by simulating the effects of fluctuations in criterion on common measures of decision-making behavior and highlight the problems that these fluctuations may cause in interpreting experimental data. We then present a novel hierarchical framework that allows trial-to-trial estimation of fluctuations in the decision-criterion and show that its parameters can be recovered even with a small number of trials (i.e., 500). Lastly, we show that this model solves the confounds caused by unaccounted fluctuations in decision criterion.

### Why should we care about fluctuations in decision criterion?

In the next section we simulate the empirical consequences of criterion fluctuations, and especially highlight how these could impact different experimental conclusions. Rather than assuming a fixed decision criterion across time, we model the criterion fluctuations as a first-order autoregressive process (AR(1); see (19) and (21) for similar approaches). An AR(1) model describes the temporal dynamics of a time series, in this case the decision criterion, via two parameters. First, an autoregressive coefficient *a* captures the temporal dependency between two successive time points (*x*_*t*_ and *x*_*t*−1_). In general, for an AR(1) model to have stable behavior, *a* must take values between -1 and +1. When *a* = 0, the resulting criterion is sampled from a normal distribution without across-trial autocorrelation. Second, a zero-mean error term with variance *σ*^2^ causes the random fluctuations. Figure 1B illustrates how variations in both these parameters influence a simulated time series of the decision criterion. Smaller magnitudes of *a* (bottom left panel) make a time series more mean-reverting, with shorter fluctuations around its mean. In contrast, when a time series is less mean-reverting due to larger values for *a* (top left panel), it will deviate more strongly from its mean for longer periods. The variance of the error term *σ*^2^ governs the scale of the random fluctuations and thus acts as a scaling parameter, as can be seen by comparing the left and right panels. The left and right panel time series show the exact same dynamics but merely differ in magnitude on the *y*-axis.

Having established a model to generate fluctuations in decision criterion, we next use simulations to show that these fluctuations can lead to biased and erroneous inferences in various commonly used metrics that (implicitly) assume stable criteria. Below, we discuss three examples highlighting the importance of accounting for a time-varying criterion. First, the presence of criterion fluctuations can mimic a causal influence of previous responses, causing apparent sequential effects in decision-making. Second, criterion fluctuations lead to biased estimates of popular measures of sensitivity. Specifically, both the slope of the psychometric function and *d*′ from SDT are underestimated in the presence of criterion fluctuations. Third, the ability to quantify criterion fluctuations from trial to trial will contribute to a better understanding of decision-making and its neural basis.

#### Criterion fluctuations can create apparent sequential effects

A wealth of studies have suggested that decisions are not isolated events, but instead show sequential effects: past trials bias the current decision. Such sequential effects have been observed in humans (22; 23; 24; 25; 26; 27; 28; 29; 30), primates (31; 20), and rodents (19; 32; 33; 34; 35) for a wide range of tasks. In 2AFC tasks, one example of a sequential effect is the tendency to repeat previous responses, also known as choice history bias, repetition bias, serial dependencies, perseveration, or hysteresis (36; 37; 38; 23; 39; 40; 28; 41; 29; 42). These effects are often taken as evidence that observers maintain active representations of their environment or decision history: an influence of previous responses is typically interpreted and modeled as causal effect, where observers update their criterion based on some combination of past stimuli, choices and rewards (22; 32; 28; 41). For example, Treisman and Williams (13) suggested that sequential effects can be explained by adjusting the criterion on the current trial as a function of the observer’s previous response.

However, sequential effects can also arise in absence of any “systematic” adjustments, purely as the consequence of random fluctuations in the decision criterion (see also 19; 32; 43). To illustrate this point, we simulate five different datasets consisting of 50 agents with 5000 trials each. Critically, we simulate behavior in which the decision criterion is not stable but instead fluctuates over time according to a first-order autoregressive process. To alter the dynamics of the criterion fluctuations, each dataset is generated with different values for *a* and *σ*^2^. On each trial, the response is drawn from a Bernoulli distribution with the probability governed by stimulus evidence (true *β*_stimulus_ = 1.25), previous response (true *β*_previous response_ = 0), and a fluctuating value over trials representing the fluctuations in criterion. Note that adding this fluctuating value to the Bernoulli probability is mathematically identical to comparing a decision variable to a varying criterion (i.e., fluctuating criterion; fig. 1A). When this value does not fluctuate (*a* = 0 and *σ*^2^ = 0) this corresponds to a fixed and unbiased criterion (i.e., static criterion; fig. 1A). In fig. 1C, we confirm that simulated observers using a fixed criterion do not display a tendency to repeat the previous response (i.e., no choice history bias). However, a fluctuating decision criterion causes the emergence of *apparent* choice history bias, despite the absence of any “systematic” criterion shifting in function of previous response (true *β*_previous response_ = 0) (see (32, figure 2), and (19) for similar simulations). As expected, this apparent choice history bias increases when the random fluctuations in decision criterion become stronger (i.e., increased *a* and *σ*^2^). In sum, researchers interested in choice history effects cannot ignore the possibility of fluctuations in the decision criterion.

**Figure 2:**
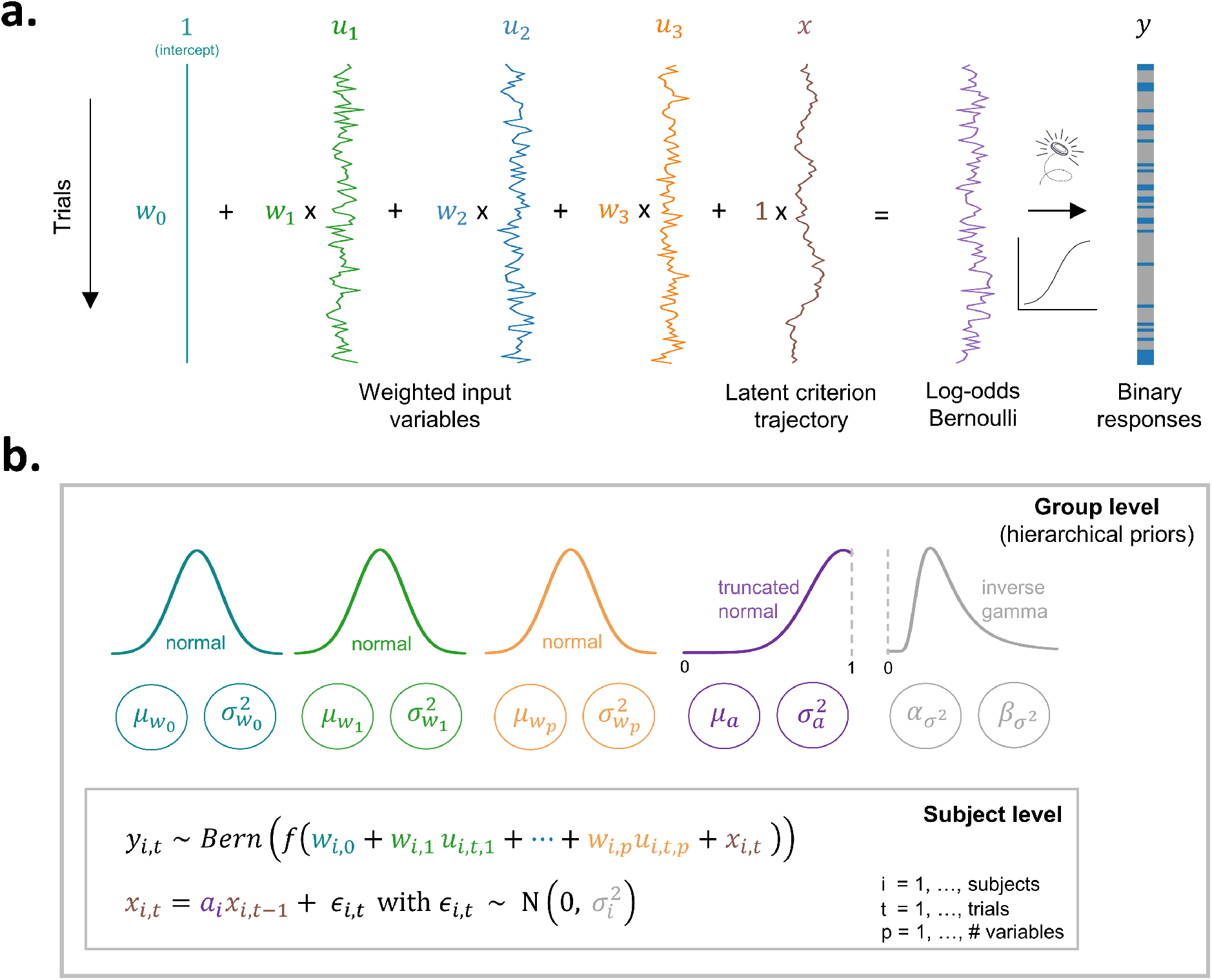
An overview of the generative model and its hierarchical structure. **A)** The figure displays, using 100 simulated trials for one agent, responses sampled from a Bernoulli distribution (i.e., a series of weighted coin flips). The trial-by-trial Bernoulli probabilities are a function of a weighted combination of the observed covariates ***u*** _*t*_ and the latent criterion fluctuations *x*_*t*_. The first column consist of a constant to allow the model to estimate an intercept *w*_0_. The next three columns show how the covariates, drawn from a standard normal distribution N(0, 1) evolve randomly over time. Summing the weighted covariates with the criterion trajectory results in the log-odds of the Bernoulli distribution. After applying a sigmoid transformation to transform log-odds into probabilities, we draw from the Bernoulli distribution (i.e., weighted coin flip) to produce binary responses. **B)** For each subject *i* we infer weights (*w*_*i*,0_, *w*_*i*,1_, …, *w*_*i,p*_) and model the latent criterion fluctuations *x*_*i,t*_ as an AR(1) process with per-subject autoregressive coefficient *a*_*i*_ and variance 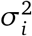. Through the specification of hierarchical priors we allow the sharing of statistical strength across subjects when estimating these individual parameters. Like the individual parameters, the parameters of the hierarchical distributions are iteratively updated during the estimation procedure.

#### Criterion fluctuations lead to an underestimation of perceptual sensitivity metrics

Many researchers are interested in perceptual sensitivity, which is often measured using one of two popular methods: the psychometric slope and *d*′ in signal detection theory (SDT). Both assess how well individuals can discriminate stimuli, and each has its own method of quantifying sensitivity. The psychometric slope is obtained by presenting stimuli at multiple levels of intensity and fitting a psychometric function to the data. The slope from this function is often directly used as a measure of sensitivity, or researchers may infer a threshold of interest by evaluating the slope at a particular stimulus intensity strength (44). However, in the presence of criterion fluctuations, the slope of the psychometric function is severely biased (fig. 1D). Specifically, the slope of the psychometric function decreases with increasing strength of criterion fluctuations, despite having the same generative psychometric slope (true *β*_stimulus_ = 1.25). This underestimation occurs because fluctuations in the criterion lead to more variable responding to identical stimuli, which ultimately flattens the psychometric function (25; 45; 14). Such distortions may impact scientific conclusions, for example when determining the threshold for conscious access (46), or a specific level of performance (47). In sum, researchers applying psychometric functions in their work should be aware of criterion fluctuations and its consequences.

Similarly, inspired by signal detection theory researchers often use *d*′, the difference between z-scored hits and z-scored false alarms, as a measure of sensitivity (45). For a given level of true sensitivity, *d*′ remains unchanged in the presence of biased decision criteria (cf. iso-sensitivity curves; (5)). Although this property holds in the presence of a fixed response bias (i.e., static criterion; fig. 1A), the property does not hold when the criterion fluctuates (11). Specifically, criterion fluctuations lead to underestimation of the true sensitivity, biasing estimates of *d*′. To demonstrate this point, we again simulate behavior for different observers that all have the same true sensitivity (i.e., same *β*_stimulus_), but differ in criterion fluctuation dynamics. The simulation follows the same procedure as before (fig. 1D) but now only includes one level of (signed) evidence. If we compute *d*′ based on the data without criterion fluctuations, we get an estimated *d*′ of 1.09 (fig. 1E). However, the presence of criterion fluctuations biases *d*′ estimates downwards. In sum, researchers interested in comparing sensitivity, as measured by *d*′, across participants or conditions should account for the possibility of criterion fluctuations.

#### Criterion fluctuations as a topic of study in human neuroscience

The above examples illustrate how criterion fluctuations can result in various biases. Note that the impact of fluctuations is not limited to only these examples: the concern that non-stationarities might underlie post-error and post-confidence slowing — i.e., slow fluctuation in the decision boundary of the drift diffusion model — has been raised in other domains as well (48; 49). While these examples stress the importance of controlling for these fluctuations, it is crucial to highlight that the criterion fluctuations are themselves an intriguing phenomenon worthy of exploration. A growing body of research illustrates that decision-making is characterized by non-stationarities (17; 18; 49; 48; 19; 50; 20; 21; 51). Whereas these non-stationarities seem to be a fundamental feature of decision-making, they are still often ignored or considered a nuisance. Recently, a general trend in cognitive neuroscience is emerging towards more dynamical approaches, with more effort spent to capture time-varying internal states (52; 16). The ability to measure trial-to-trial criterion fluctuations opens up new avenues for research, such as uncovering the neural mechanisms underpinning them, identifying more phenomena in which they play a role, and leveraging this understanding to improve computational models. For example, quantifying criterion fluctuations with an AR(1) model and estimating its parameters allows the comparison of these parameters over subjects and experimental manipulations. In sum, studying criterion fluctuations will advance our understanding of decision-making and its underlying mechanisms.

### Previous attempts to measure criterion fluctuations

In the previous section, we showcase different scenarios illustrating the importance of considering fluctuations in the decision criterion. In the context of choice history bias (i.e., repetition effects in function of previous response), earlier work attempted to control for criterion fluctuations through a model-free approach (32; 43), similar to one developed in the post-error slowing literature (48). Since the criterion is assumed to be autocorrelated, it should have a similar influence on temporally adjacent trials. In theory, it should thus be possible to dissociate: (i) a causal influence of the previous response on the current trial, from (ii) an acausal influence of the next response on the current trial, which reflects the value of the criterion that varies only slowly across several trials. Lak et al. (32) proposed to obtain a drift-free measure of choice history bias by subtracting the acausal from the causal influence. Although this model-free approach sounds promising, Gupta and Brody (19) demonstrated that this is only effective in the context of a specific subset of updating strategies (i.e., no systematic updating and symmetric win-stay lose-switch). In the presence of other updating strategies, such as win-stay lose-stay or win-stay lose-nothing, their model-free correction resulted in inaccurate and biased estimates. Therefore, this proposed model-free approach is not an appropriate tool to correct for criterion fluctuations. Moreover, it does not allow extracting a trial-by-trial measure of the latent criterion to study its neural and physiological basis.

Instead of the model-free approach, Gupta and Brody (19) have proposed a model-based approach which explicitly models criterion fluctuations as an AR(1) process in a Bernoulli linear dynamical system. With this approach they were able to correctly recover a wide range of updating strategies. However, there are some key limitations that prevent a widespread usage of their approach, especially in human cognitive neuroscience. First, the authors only tested their model using very large datasets: simulated datasets with 40.000 trials, or rat data with 50.000 trials, obtained over many sessions. Second, their model exhibited inherent trade-offs between parameters. Specifically, their model featured a parameter that weighted the criterion fluctuations in the Bernoulli probability. This generated a trade-off with the error variance (*σ*^2^) as both control the scaling of the criterion fluctuations. Finally, the authors only focused on the recovery of history biases but did not quantitatively investigate the recovery of the criterion trajectory and its underlying computational parameters. Instead, they assumed a fixed autoregressive coefficient a of .9995. Similarly, Roy et al. (21) developed a method to estimate trial-by-trial fluctuations in the weights of a logistic regression model, which captures criterion fluctuations through the intercept. However, the weights were assumed to evolve over time according to an AR(1) process with *a* fixed to 1 (i.e., a random walk). Given that the parameter *a* plays a crucial role in capturing the temporal dynamics of a time series (i.e., as shown in fig. 1B), fixing this parameter to a specific value is therefore an overly strong assumption. It is likely that the generative autoregressive coefficient of criterion fluctuations in experimental data deviates from this value, which may bias estimates of trial-to-trial criterion trajectories.

## Hierarchical Model for Fluctuations in Criterion (hMFC)

In the current work, we develop the Hierarchical Model for Fluctuations in Criterion (hMFC), which aims to solve these limitations by extending earlier approaches in various respects. First, to solve the issue that the model requires a high number of trials to be fitted, we implement a Bayesian hierarchical estimation approach. This allows to combine data across multiple participants and concurrently models both group level estimates and individual estimates. Therefore, fewer trials are required to obtain reliable parameter estimates. Second, we adjust our model structure compared to Gupta and Brody (19) to avoid the parameter trade-off discussed earlier. Specifically, in hMFC this trade-off is solved by not additionally weighing the criterion fluctuations in the Bernoulli probability (i.e., setting it to 1). Third, instead of fixing *a* in the AR(1) to .9995 (19) or 1 (21), we implement this in our model as a free parameter. This allows to capture a wide range of criterion fluctuation dynamics and ultimately enables researchers not just to control for criterion fluctuations, but also to study these fluctuations and its dynamics as the main topic of investigation (e.g., comparing a over different experimental manipulations or investigating individual differences). Finally, we provide easy to use code and software demos to enable the application of our model by the cognitive neuroscience community.

In sum, in order to model criterion fluctuations in decision criterion we implement a hierarchical framework within a Bernoulli linear dynamical system, called the Hierarchical Model for Fluctuations in Criterion (hMFC). Next, we elaborate on the technical details of hMFC.

### The generative model

First, consider a single session consisting of a sequence of *T* trials. On each trial *t* = 1, …, *T*, the agent makes a binary response *y*_*t*_. Our generative model assumes the response is drawn from a Bernoulli distribution:

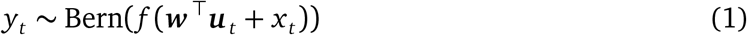

where the probability of a response is governed by the covariates ***u*** _*t*_ ∈ ℝ^*p*^, the latent state *x*_*t*_ ∈ ℝ, and the weights ***w*** ∈ ℝ^*p*^. The sigmoid transformation, *f* (*z*) = 1/(1 + *e*^−*z*^), constrains the conditional probability to be between 0 and 1. The vector ***u*** _*t*_ contains a constant of 1 for the model to estimate an intercept, and observed input variables or covariates on trial *t*, such as the stimulus evidence or previous response. The influence of these covariates on the Bernoulli probability is captured by the weights ***w*** ∈ ℝ^*p*^. The latent state *x*_*t*_ ∈ ℝ on trial *t*, which is not directly observable, is assumed to follow a first-order autoregressive process:

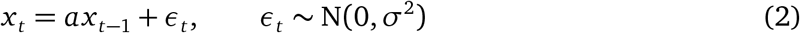

where *a* ∈ [0, 1] is the autoregressive coefficient which captures the temporal dynamics of the criterion fluctuations (fig. 1B), and *σ*^2^ controls the variance of the fluctuations (i.e., *σ*^2^ is a scaling parameter). Note that adding a drifting value *x*_*t*_ to the log-odds of the Bernoulli distribution is mathematically equivalent to comparing a decision variable to a fluctuating criterion (fig. 1A). In sum, the model predicts responses based on a combination of the weighted variables *u*_*t*_ and the latent state *x*_*t*_, as illustrated in fig. 2A. Ultimately, this model allows us to disentangle two sources that influence the response probability: (i) the influence of ***u*** _*t*_, and (ii) the influence of the latent state *x*_*t*_.

### The hierarchical prior: putting the ‘h’ in hMFC

Having explained the generative model and the behavior it generates (fig. 2), we next focus on the inverse step, namely estimating the underlying latent parameters given observed behavior. In many settings, we need to obtain accurate parameter estimates from only tens to hundreds of trials per subject. Small values of *T* pose statistical challenges for parameter estimation. However, we can pool information across individuals to aid in inference. To this end, we developed a hierarchical Bayesian model that allows for per-subject parameter estimates, while sharing information across subjects via a global prior distribution. We infer the per-subject parameters, the latent states (i.e., criterion fluctuations) for each subject, as well as the strength of the hierarchical prior using a fully Bayesian inference algorithm.

In the hierarchical model, each subject *i* = 1, …, *N* has its own responses *y*_*i,t*_, covariates *u*_*i,t*_, and latent states, *x*_*i,t*_, for *t* = 1,…, *T*_*i*_. Each subject also has its own parameters *θ*_*i*_ = (***w*** _*i*_, *a*_*i*_, 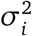), where ***w*** _*i*_ ∈ ℝ^*p*^ denote the covariate weights, *a*_*i*_ ∈ [0, 1] is the autoregressive parameter, and 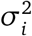 is the variance of the latent criterion fluctuations. We learn a hierarchical prior that captures how the subject-level parameters *θ*_*i*_ vary across the population. The hierarchical prior allows us to share statistical power from other subjects when estimating *θ*_*i*_. This results in better parameter recovery, even when a relatively small number trials per subject is available (53).

We specified hierarchical priors for the three types of per-subject parameters. First, we assume a Gaussian prior on the weights,

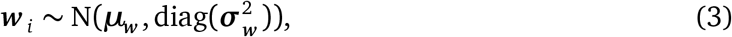

where ***μ***_***w***_ ∈ ℝ^*p*^ and 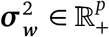 denote the global mean and variance of the per-subject weights. Second, we assume a truncated normal prior on the per-subject autoregressive coefficient,

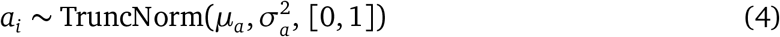

where *μ*_*a*_ and 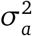 are the global mean and variance, respectively, and the domain is constrained to [0, 1] because negative values of *a*_*i*_ would lead to large jumps in the criterion, and values greater than 1 would lead to unstable dynamics. Third, we specify an inverse gamma (IGa) prior for the variance of the criterion fluctuations,

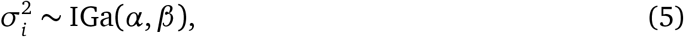

with shape parameter *α* ∈ ℝ_+_ and a scale parameter *β* ∈ ℝ_+_.

Finally, we place weakly informative priors on the parameters of the hierarchical model, *η* = (***μ***_***w***_, 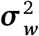, *μ*_*a*_, 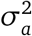, *α, β*), see the Appendix for complete details. These global parameters will be inferred and iteratively updated alongside the per-subject parameters using a fully Bayesian inference algorithm, which we describe next.

### A posterior inference algorithm

We develop a fully Bayesian inference algorithm to estimate the the posterior distributions of the per-subject latent states and parameters, as well as the parameters of the hierarchical prior. We use an augmented, blocked Gibbs sampling algorithm to produce samples that are asymptotically drawn from the desired posterior distribution.

The Gibbs sampling algorithm proceeds by iteratively sampling the conditional distribution of one block of variables, holding the rest fixed. However, this approach requires the closed-form expressions for the conditional distributions. This tractability can be achieved if the model is conditionally conjugate. Unfortunately, the Bernoulli likelihood is not conjugate with the Gaussian hierarchical priors for the covariate weights or the AR(1) model for the latent states. To circumvent this challenge, we employ the Pólya-gamma augmentation trick (54). With this augmentation, we can obtain a closed-form expression for the per-subject covariate weights ***w*** _*i*_ given the response, covariates, criterion fluctuations, and the hierarchical prior.

For the two other per-subject parameters, *a*_*i*_ and 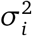, closed-form expressions for the conditional posteriors are available. First, for *a*_*i*_ the truncated normal prior is conjugate with the AR(1) model likelihood. Similarly, the inverse gamma prior for 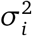 is conjugate with the AR(1) model of the latent criterion fluctuations.

After Pólya-gamma augmentation, the conditional distribution of the latent criterion fluctuations is a chain-structured Gaussian graphical model. We generate exact samples from this conditional distribution using a forward filtering/backward sampling algorithm (55). The forward filtering step computes the filtering distributions of the latent criterion fluctuations, given all the responses up to a certain trial. Once the filtering distributions have been computed for each trial, the backward sampling step generates a sequence of latent fluctuations, working backward from the last trial *T*_*i*_ down to the first. This procedure produces a sequence of latent fluctuations 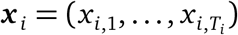 that is distributed according to its conditional distribution given the per-subject parameters, covariates, responses, and Pólya-gamma augmentation variables. Note that after estimating the latent states, we mean-center the states to ensure that the per-subject intercept *w*_*i*,0_ remains identifiable.

Finally, we update the parameters of the hierarchical prior *η* by sampling from their conditional distribution under a weakly informative prior. For the two group-level parameters of the normal priors on the covariate weights, ***μ***_***w***_ and 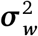, we have closed-form expressions for the conditional distributions. We update the remaining hierarchical prior parameters — *μ*_*a*_, 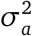, *α* and *β* — using random-walk Metropolis-Hastings with a symmetric Gaussian proposal distribution.

Together, these updates yield a Markov chain whose stationary distribution is the posterior distribution over the latent per-subject fluctuations, per-subject parameters, and hierarchical prior parameters. Repeatedly applying these updates yields samples that are asymptotically distributed according to this posterior. We can use these samples to estimate posterior expectations and perform hypothesis tests, as demonstrated in the results below. Complete details of the augmented, blocked Gibbs sampling algorithm can be found in the Appendix.

### Accessing the model

Our modeling and inference code, together with a comprehensive demonstration, are available at www.github.com/robinvloeberghs/hMFC. In addition, more in-depth plots for the simulation results are also available at this link.

## Results

To assess the effectiveness of the hMFC parameter estimation, we simulate data with a varying number of trials (500, 1000, 2500 or 5000 trials) per subject, with 50 subjects per dataset and 50 datasets per trial count. For each subject *i*, four weights, ***w*** _*i*_ = (*w*_*i*,0_, …, *w*_*i*,3_), are independently drawn from a normal distribution with mean ***μ***_***w***_ = (0.0, 0.2, −0.3, 0.6) and variance 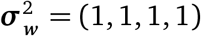. Similarly, the covariates ***u*** _*t*_ ∈ ℝ^4^ are also sampled from a standard normal distribution (mean zero, identity covariance), except for *u*_*i*,0_, which is a constant of 1 to allow the model to estimate an intercept (*w*_*i*,0_). The criterion fluctuations are sampled from an AR(1) model with the autoregressive coefficient *a* drawn from a truncated normal (mean 0.99, standard deviation 0.025, minimum value 0, and maximum value 1) and 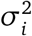 sampled from an inverse gamma distribution (*α* = 5.0, *β* = 0.45).

### Group-level (global) parameter recovery

The group-level (global) parameters *η* = (***μ***_***w***_, 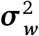, *μ*_*a*_, 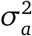, *α, β*) are accurately recovered. For almost all datasets, the generative value lies within the 95% credible interval of the posterior estimates. Most important, for ***μ***_***w***_ and 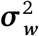 the 95% credible interval contains the true value for all datasets. Note that the width of these posteriors stays relatively constant for different trial counts (fig. S2). On the other hand, the posterior width does go down with an increasing number of subjects. In Figure 3 it is shown that when increasing the number of subjects while keeping the number of trials constant (n=500), the width of the posteriors becomes smaller. Jointly, this suggests that in order to make more reliable inferences at the group level, for hMFC it is more advisable to gather more subjects rather then gathering more trials per subject.

**Figure 3:**
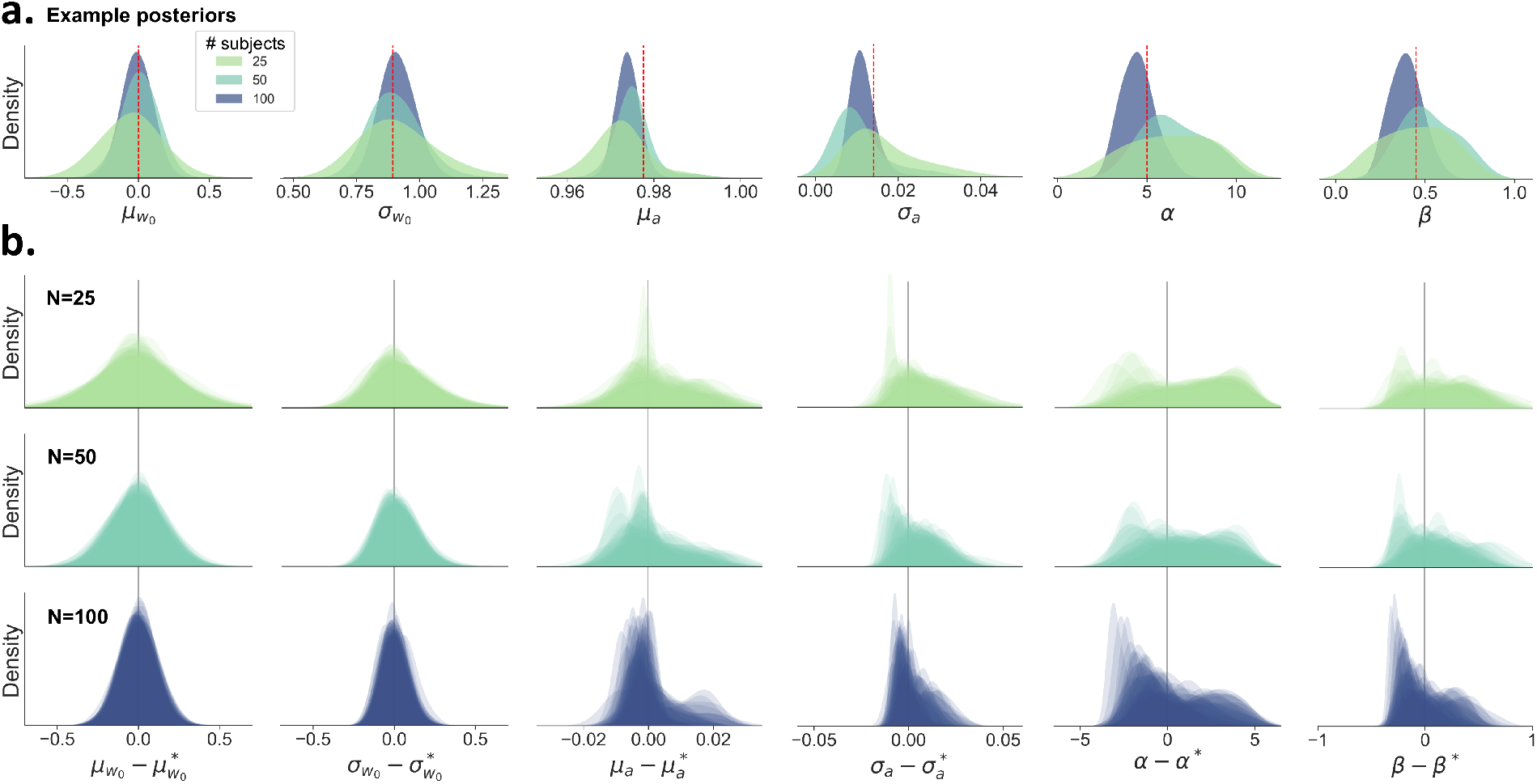
An overview of the estimated posteriors distributions for the group-level (global) parameters when varying the number of subjects per dataset, with 500 trials per subject. **A)** The posterior distributions become more narrow as the number of subjects increases. The datasets were chosen to have matched true parameter values, indicated by the red dashed line. **B)** Each row shows the overlaid posteriors for all 50 simulated datasets with a varying number of subjects per dataset. The estimated posteriors are corrected and centered on the true value (denoted by an asterisk). For 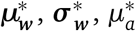, and 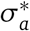, the true value is defined as the mean or standard deviation of the true per-subject parameters ***w*** _*i*_ and *a*_*i*_ within each dataset. Note that the posteriors can be skewed due to boundaries in the parameter space. Specifically, 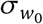, *σ*_*a*_, *α*, and *β* cannot become negative, resulting in right-skewed posteriors. Also *μ*_*a*_ is restricted in that it cannot exceed 1, as this would result in unstable behavior. Overall, we see an excellent recovery of the group-level (global) parameters.

### Per-subject (local) parameter recovery

The inference algorithm can generally recover all per-subject parameters (Figure 4A). The per-subject weights *w*_*i*,0_, *w*_*i*,1_, *w*_*i*,2_, *w*_*i*,3_ show excellent recovery (*r >* .98), even for datasets with only 500 trials per subject (Figure 4D). The autoregressive model parameters *a*_*i*_ and 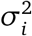, which capture the dynamics of the criterion fluctuations, show an excellent recovery with many trials per subject (*r* = .95 for *a*_*i*_, *r* = .89 for 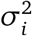, with 5000 trials), but are less accurately estimated when fewer trials per subject are available (*r* = .66 for *a*_*i*_ and *r* = .56 for 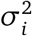 with 500 trials) (Figure 4B,C).

**Figure 4:**
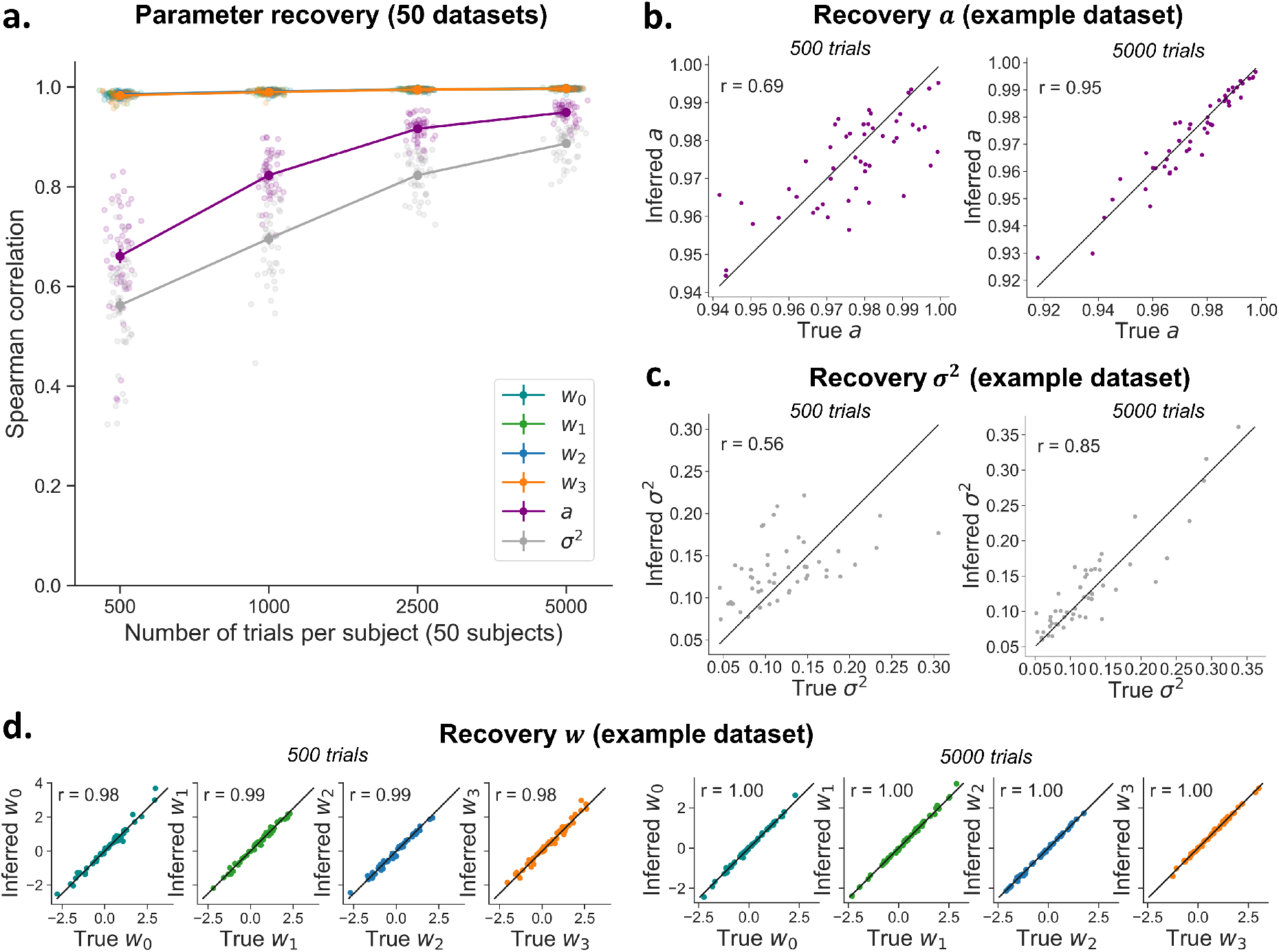
Recovery of the per-subject parameters of the hMFC model. **A)** The covariate weights *w* show excellent recovery even with low trial numbers (note that the lines for the different weights overlap). The model is able to recover *a* and *σ*^2^ very well with high trial numbers, but recovery drops with a limited number of data points per subject. Each dot represents the recovery correlation for one dataset. Note that error bars (standard error) are shown but they are very small. **B-D)** The recovery for *a* (B) and *σ*^2^ (C) and covariate weights *w*_0_, *w*_1_, *w*_2_, *w*_3_ (D) is shown for an example dataset with 500 trials and 5000 trials per subject. Each dot represents one subject. The line shows the diagonal.

### Per-subject criterion trajectory recovery

Importantly, although the per-subject parameters *a*_*i*_ and 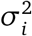 show somewhat low recovery with low trial numbers, the model can still very accurately recover the trial-to-trial latent criterion trajectory (Figure 5) crucial for investigations into its neural and physiological bases. The recovery, measured as the correlations between true and inferred latent trajectory, is generally high (*r* > .80), and consistent across datasets (fig. 5A). As shown for one example subject, the true and the inferred trajectory strongly correlate (*r* = .85), even though the model is not able to perfectly recover high-frequency fluctuations in the true latent trajectory (fig. 5C).

**Figure 5:**
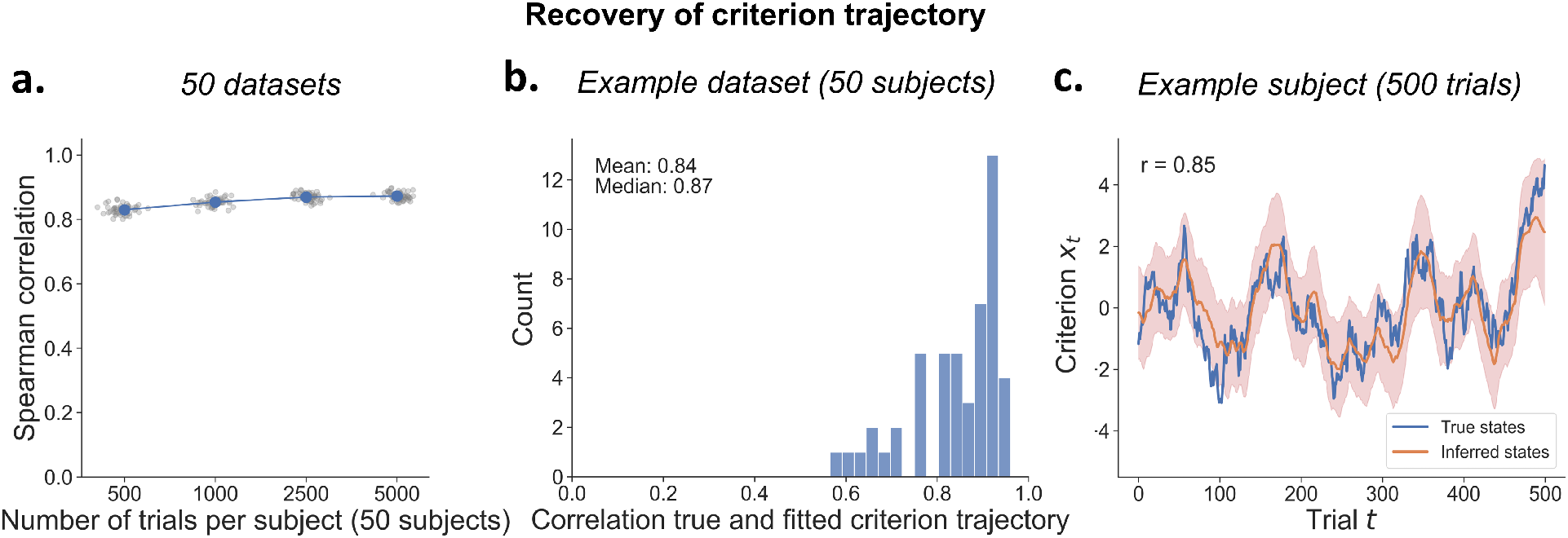
Recovery of the latent criterion trajectory. **A)** Each dot represents the criterion trajectory of one dataset, averaged over subjects. Note that error bars (standard error) are plotted but are very small. **B)** A representative dataset of 50 subjects with 500 trials each shows an average correlation between true and inferred criterion fluctuations of *r* = .84. **C)** True and inferred criterion fluctuations for an example subject. The shaded area represents the 95% credible interval of the estimated posterior at each trial.

At lower trial counts, recovery ranges from 0.57 to 0.96. across subjects (fig. 5B). This variation is partly due to the stochastic criterion fluctuations themselves, but also relates to the combination of parameters governing the individual AR(1) processes. Specifically, trajectory recovery is more difficult for lower values of *a* combined with higher values of *σ*^2^ (fig. S3).

### Computation time

Fitting the data of 50 subjects with 500 trials each for 1000 iterations takes around 1 hour on a regular laptop with only CPU’s. When GPU’s are available the computation time is reduced to 5 minutes, making this procedure sufficiently lightweight to run easily on typical human datasets even for researchers without dedicated computing facilities.

### Application hMFC to sequential effects and measures of sensitivity

Having verified that the inference procedure recovers the generative hMFC model, we assess if it also corrects for the spurious effects that can arise from ignoring criterion fluctuations: apparent sequential effects and underestimated *d*′ (as shown in fig. 1). We fit the hMFC model with stimulus evidence and previous response as covariates, to the data shown in fig. 1. Importantly, we ran the fits either with or without estimating criterion fluctuations, allowing us to test if the inclusion of these parameters accurately solves the confounds.

When criterion fluctuations are not estimated, a clear effect of previous response (i.e., tendency to repeat previous response) artificially emerges in the model (fig. 6A): the posterior distributions are shifted further away from zero when the criterion fluctuations become stronger (i.e., replicating the finding from fig. 1C, using our model fit). Fortunately, when criterion fluctuations are estimated, this estimation bias disappears entirely and posteriors center around the generative value (*β*_previous response_ = 0, red line). This shows that our model can accurately distinguish sequential effects caused by slow fluctuations in the decision criterion from sequential effects caused by systematic updating.

**Figure 6:**
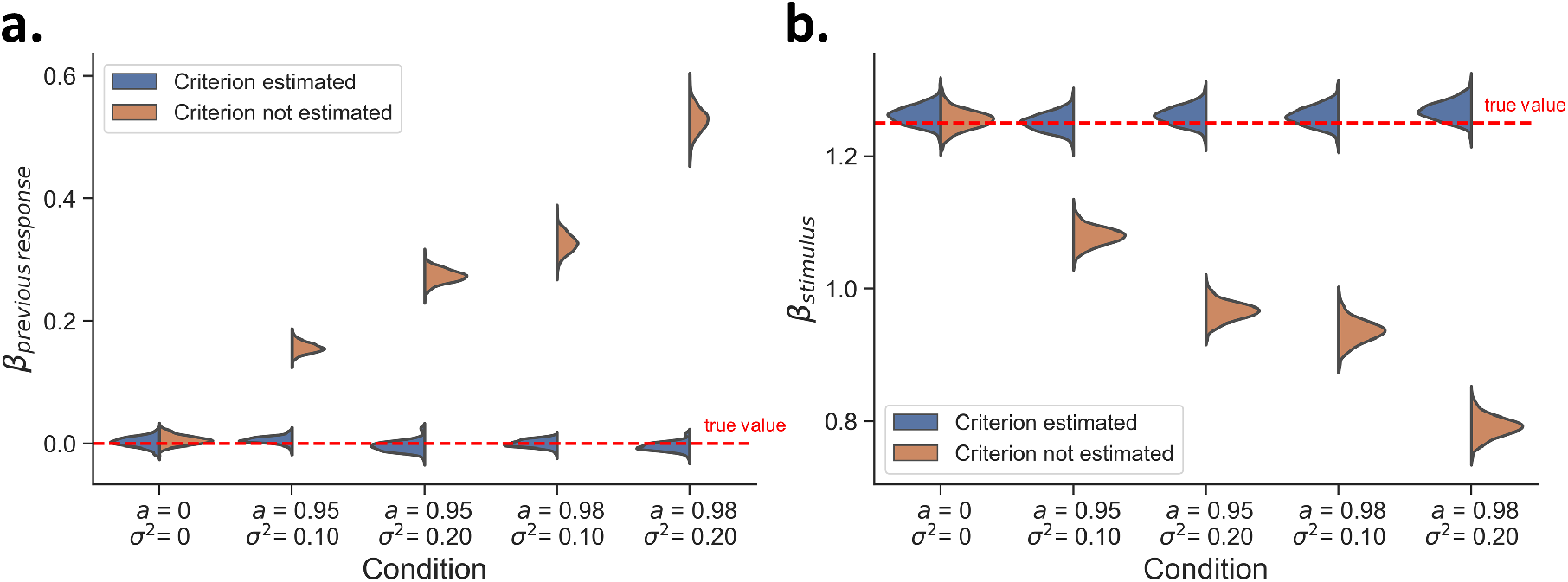
Using hMFC to account for criterion fluctuations. The posterior distributions for *β*_stimulus_ and *β*_previous response_ are shown after fitting hMFC to the simulated datasets used in fig. 1C and fig. 1D. **A)** The estimation of criterion fluctuations allows hMFC to correctly recover the absence of any systematic criterion updating in function of the response on the previous trial. **B)** Estimating criterion fluctuations helps to address the issue of underestimating the psychometric slope.

In a similar vein, when criterion fluctuations are not estimated, the psychometric slopes are systematically underestimated (fig. 6B) compared to the generative value (*β*_stimulus_ = 1.25, red line). Fortunately, estimating criterion fluctuations resolves this underestimation. In sum, the hMFC corrects for estimation biases by explicitly estimating the fluctuations in the criterion.

## Discussion

Computational models of decision-making typically assume stationarity in the parameters underlying the decision computation. However, an increasing body of research highlights the dynamic nature of decision making and shows that computational variables underlying decision-making may fluctuate over time. One example of these non-stationarities are fluctuations in the decision criterion. Ignoring such latent fluctuations can lead to false interpretations and inaccurate estimates. Specifically, criterion fluctuations can lead to behavior that mimics a causal relationship in sequential effects, and they can lead to an underestimation of two popular measures of perceptual sensitivity (*d*′ in SDT and the psychometric slope).

Here, we developed the Hierarchical Model for Fluctuations in Criterion (hMFC), a Bayesian hierarchical model to obtain trial-by-trial estimates of the decision criterion. Even with only 500 trials per participant, which is common in studies on human cognitive neuroscience, the model parameters and the latent criterion trajectory itself can be accurately inferred. Importantly, the hMFC can effectively correct for the biases that occur when criterion fluctuations are ignored.

We provide open access to the code for hMFC and created an interactive demo for researchers interested in applying this model to their data. Through this work, we hope to enable further research in human cognitive neuroscience aimed at understanding the dynamics of decision-making.

## Acknowledgements

AEU is supported by a Veni fellowship from the Netherlands Organisation for Scientific Research. SWL is supported by grants from the NIH BRAIN Initiative (U19NS113201, R01NS131987, R01NS113119, & RF1MH133778), the NSF/NIH CRCNS Program (R01NS130789), and the Sloan, Simons, and McKnight Foundations.

## Appendix

### Hierarchical Model for Fluctuations in Criterion (hMFC) Details

**Notation** Let

- *N* denote the **number of subjects**
- *T*_*i*_ denote the **number of trials** for subject *i*
- *η* = (***μ***_***w***_, 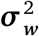, *μ*_*a*_, 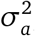, *α, β*) denote the **group-level (global) parameters**.
- *θ*_*i*_ = (*a*_*i*_, ***w*** _*i*_, 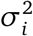) denote the **subject-level (local) parameters** for subject *i*
- *x*_*i,t*_ ∈ ℝ for *t* ∈ 1, …, *T*_*i*_ denote the sequence of **latent states** for subject *i*.
- ***u*** _*i,t*_ ∈ ℝ^*p*^ for *t* ∈ 1, …, *T*_*i*_ denote the sequence of **inputs** for subject *i*.
- *y*_*i,t*_ ∈ {0, 1} for *t* ∈ 1, …, *T*_*i*_ denote the sequence of **binary observations** for subject *i*.

The Hierarchical Model for Fluctuations in Criterion (hMFC) defines a joint distribution over latent states, observations, and parameters given the inputs,

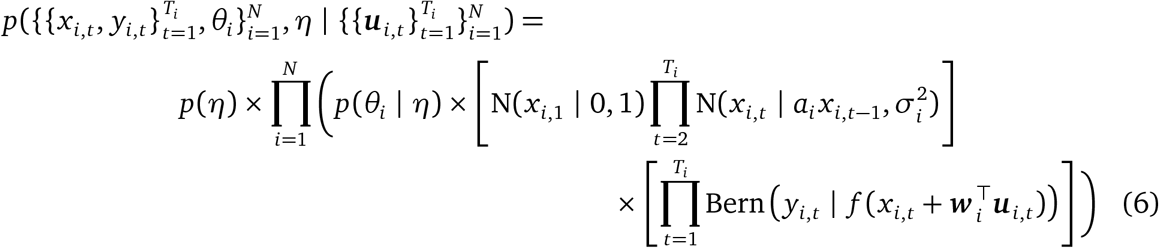

where *f* (*z*) = 1*/*(1 + *e*^−*z*^) denotes the logistic function and,

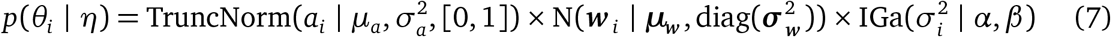

is the hierarchical prior. The truncated normal distribution ensures that *a*_*i*_ ∈ [0, 1], so that the latent state dynamics for subject *i* are stable. The hierarchical Gaussian prior on weights ***w*** _*i*_ ∈ ℝ^*p*^ allows for sharing of statistical strength across subjects.

We assume weakly informative priors over the global parameters,

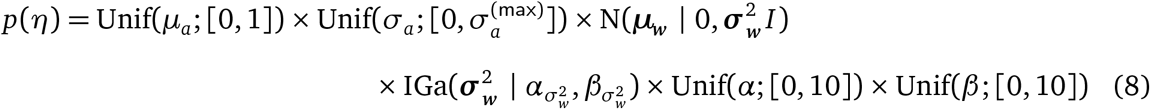

with 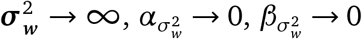. The uniform priors for *μ*_*a*_, *α*, and *β* allow for a wide range of global parameters, while still enforcing marginal stability constraints.

We set 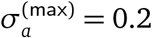 to penalize the model for using very different values for *a*_*i*_ across subjects. Nevertheless, this value is a generous upper bound.

### Posterior Inference

We designed an augmented, blocked Gibbs sampling algorithm to estimate the posterior distribution, 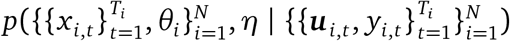.

#### Gibbs sampling the latent states

The trick is to represent the Bernoulli likelihood as a scale-mean mixture of Gaussians using the Pólya-gamma (PG) augmentation (54),

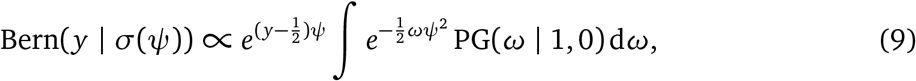

where PG(*ω* | 1, 0) denotes the standard Pólya-gamma density on *ω* ∈ ℝ_+_. In our case, 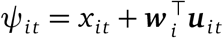. After augmenting the model with auxiliary variables *ω*_*it*_, the log likelihood as a function of *x*_*it*_ is,

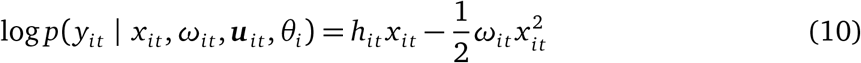

where

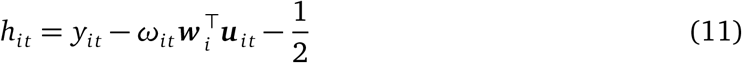

is the precision-weighted mean, which we recognize as a kind of residual.

After augmentation, the conditional probability of *x*_*i*_ is an exponentiated quadratic,

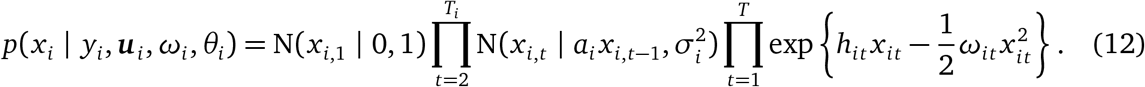

We recognize this as a chain-structured Gaussian graphical model — i.e., a linear Gaussian dynamical system — over *x*_*i*_. We use information-form message passing algorithms implemented in Dynamax to sample the latent states from their conditional distribution given the emissions, inputs, parameters, and auxiliary PG variables.

#### Sampling the Pólya-gamma augmentation variables

The conditional distribution over auxiliary variables *ω*_*it*_ given the states and parameters is,

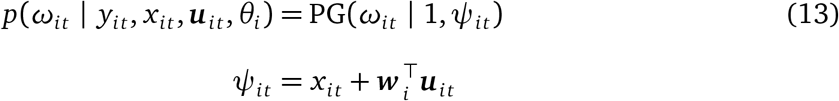

There are highly efficient rejection sampling algorithms for the PG(1, *Ψ*) distribution (56), but we can also use a naïve sampling algorithm based on a representation of the PG distribution as a weighted sum of gamma random variates (54).

Note that the augmentation variables (one per subject and trial) are conditionally independent of one another and thus can be sampled in parallel. Our implementation using the JAX library (57) takes advantage of this opportunity for fast parallel sampling on GPUs.

#### Sampling the per-subject (local) parameters

##### Input weights

After augmentation, log likelihood as a function of the weights ***w*** _*i*_ for subject *i* is,

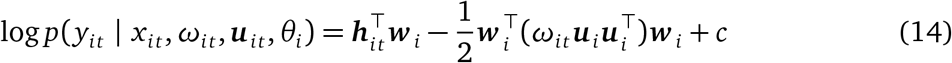

where 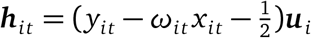 and *c* is constant w.r.t. ***w*** _*i*_.

Again, this quadratic log probability is conditionally conjugate with the Gaussian prior on the weights,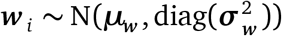. The conditional posterior distribution is,

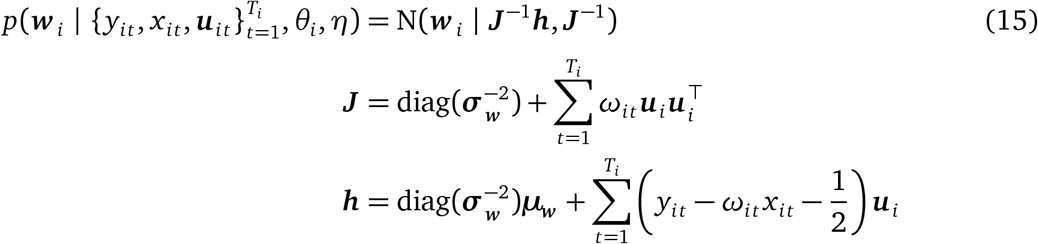

##### Dynamics coefficient

The truncated normal prior on *a*_*i*_ is conjugate with the linear Gaussian dynamics model. Given the latent states,

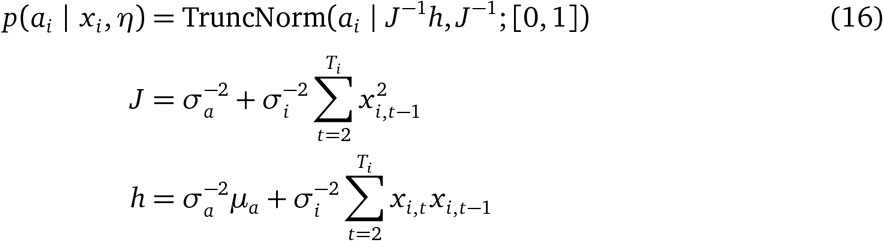

This is a truncated normal distribution with mean *J*^−1^*h* and variance *J*^−1^, constrained to *a*_*i*_ ∈ [0, 1].

##### Dynamics noise variance

The inverse gamma prior is conjugate with the Gaussian noise model. The conditional posterior distribution is,

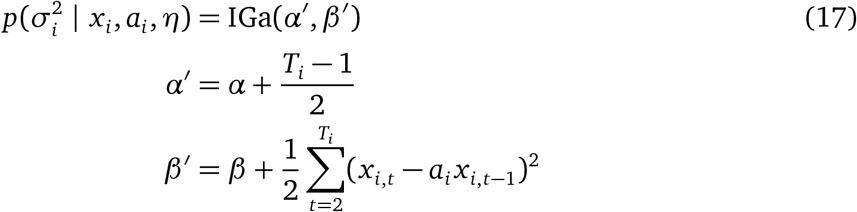

###### Sampling the group-level (global) parameters

The global parameters ***μ***_***w***_ and 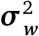 admit closed form Gibbs updates. Under their uninformative priors,

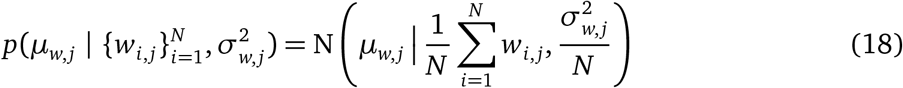

and

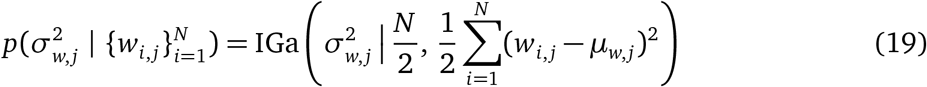

for *j* = 1, …, *p* with *p* the total number of covariates. Finally, we update the global parameters *μ*_*a*_, 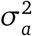, *α*, and *β* using random-walk Metropolis-Hastings with symmetric Gaussian proposal distributions.

###### Initialization

We initialize the global parameters conservatively, setting *μ*_*a*_ = 0.95, *σ*_*a*_ = 0.1, ***μ***_***w***_ = (0, …, 0)^⊤^, *σ*_*w, j*_ = 1.25 for all *j, α* = 4, and *β* = 4 × .4^2^. Thus, at initialization, the global prior is biased toward producing per-subject latent states that are reasonably autocorrelated. Then we sample the parameters and states from their conditional distributions given the global prior, without conditioning on the data.

###### Algorithm 1: Gibbs sampling algorithm for the hierarchical Bernoulli LDS

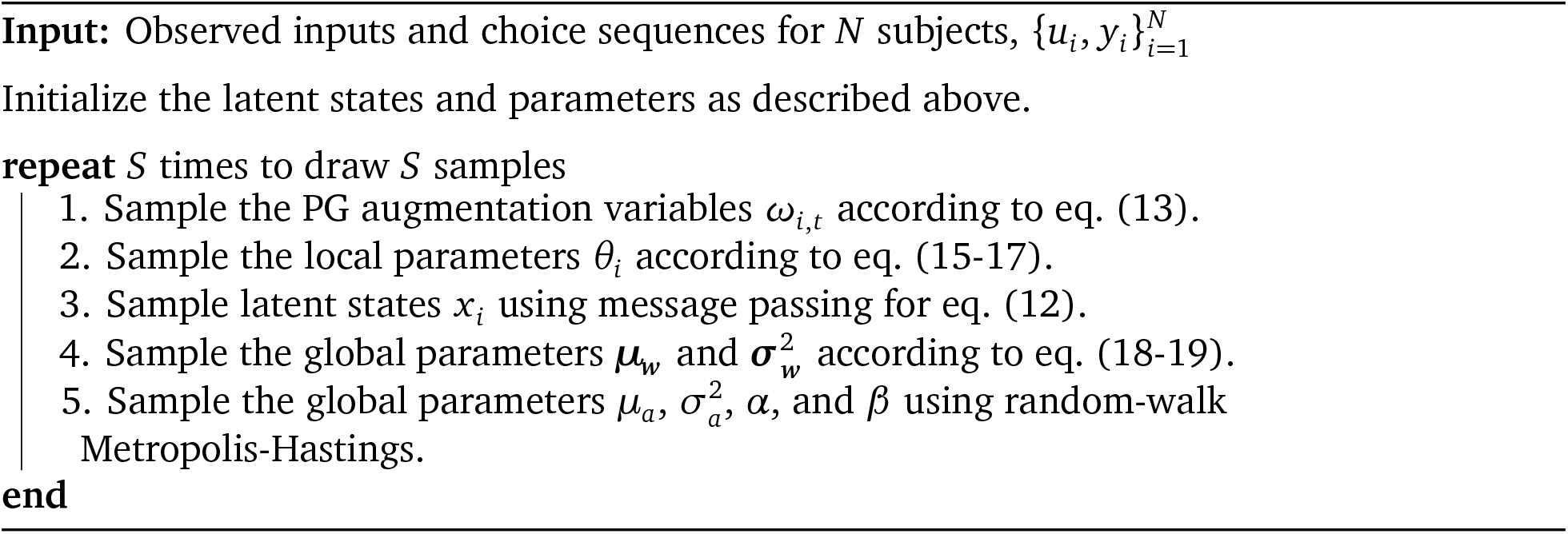

More informative initializations are possible. For example, one could fit a logistic regression (without time-varying latent states) for each subject to obtain initial weights ***w*** _*i*_, and then initialize the per-subject latent states to *x*_*i*_ = (0,…, 0)^⊤^. Based on the initial weights, we can estimate the global weight mean ***μ***_***w***_ and variance 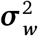. Combined with the initializations for *μ*_*a*_, 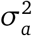, *α*, and *β* above, this procedure should effectively start the Gibbs sampler near a reasonable basin of the posterior.

###### Full algorithm

The complete algorithm is given by Algorithm 1.

## Supplemental Figures

**Figure S1:**
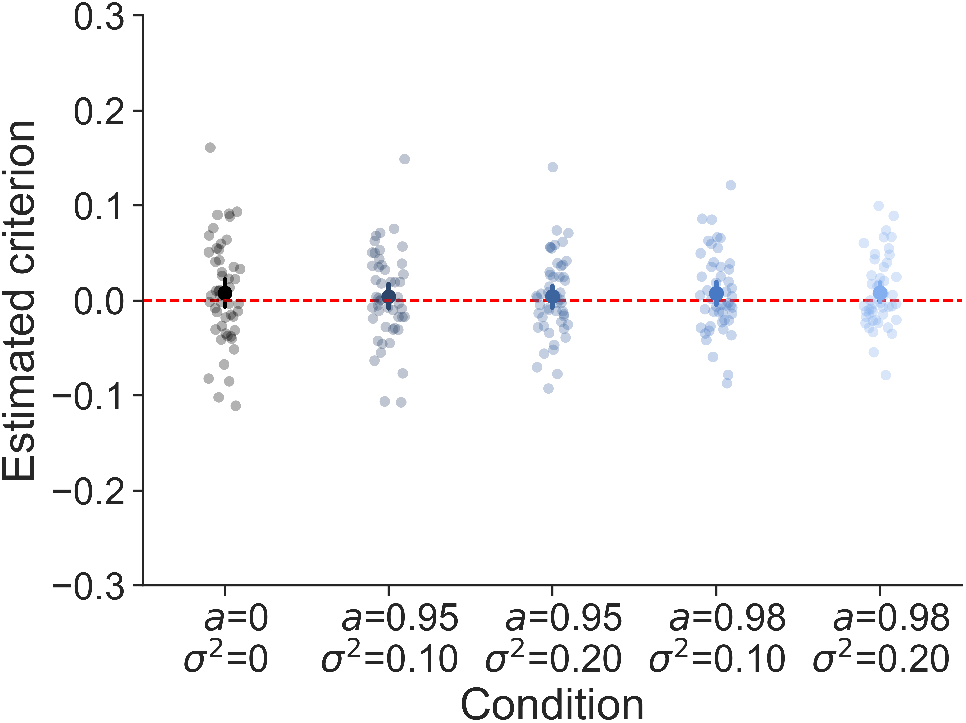
Whereas the presence of criterion fluctuations leads to an underestimation of *d*′, a popular signal-detection measure of sensitivity (fig. 1E), the criterion is on average correctly estimated when using the traditional method assuming a stable criterion. The red line shows the true criterion value.

**Figure S2:**
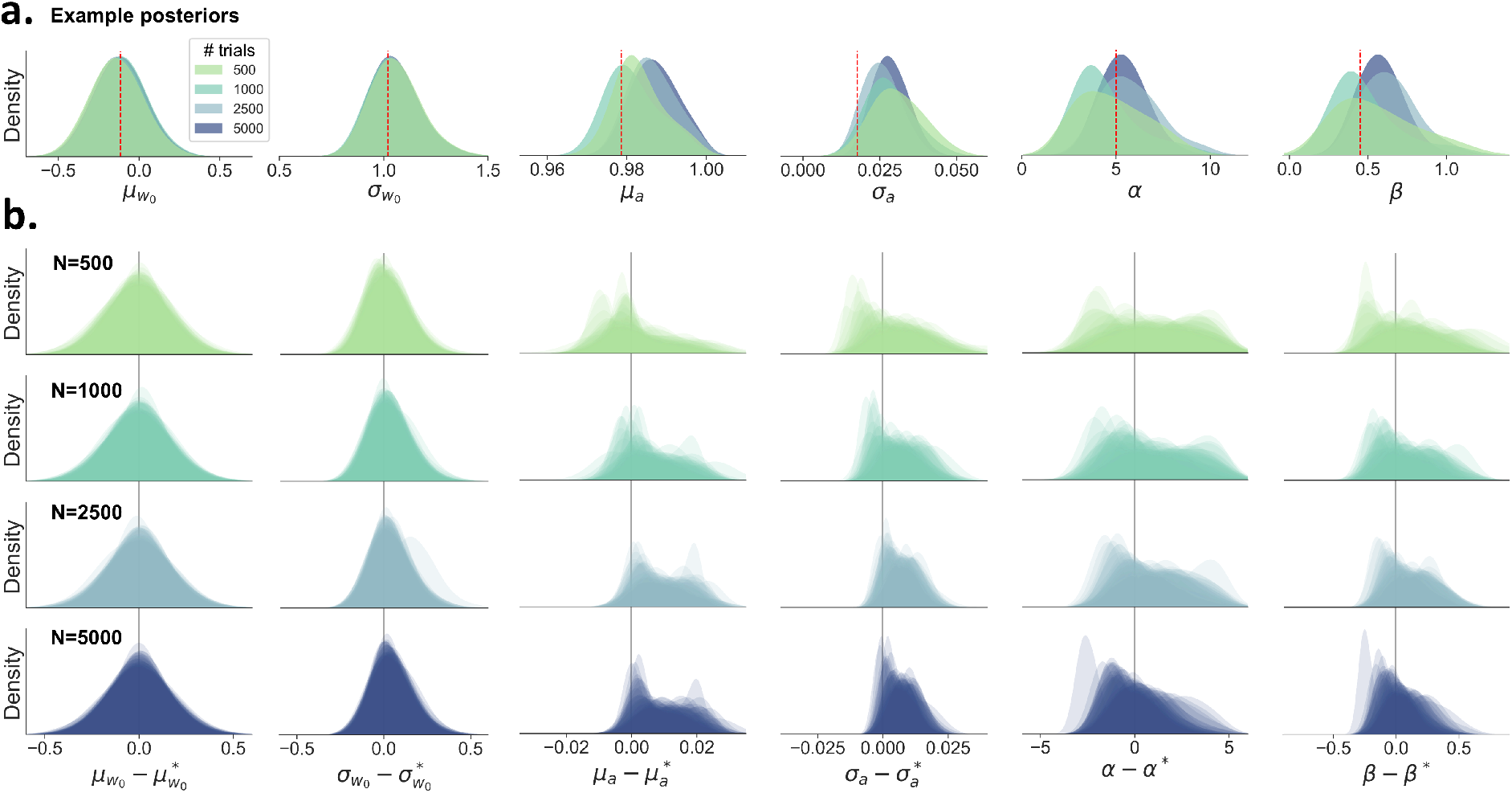
The recovery of the group-level (global) parameters in function of the number of trials per subject (500, 1000, 2500, 5000). In total, 50 datasets were simulated, each with 50 subjects. A) An example posterior is shown for each parameter and subject level. In contrast to fig. 3, where the posteriors become more narrow when increasing the number of subjects, here we see that the posterior precision remains the same with increasing trial counts. B) The overlaid posteriors are shown for all 50 datasets. The estimated posteriors are corrected and centered on the true value (denoted by an asterisk), similar to fig. 3.

**Figure S3:**
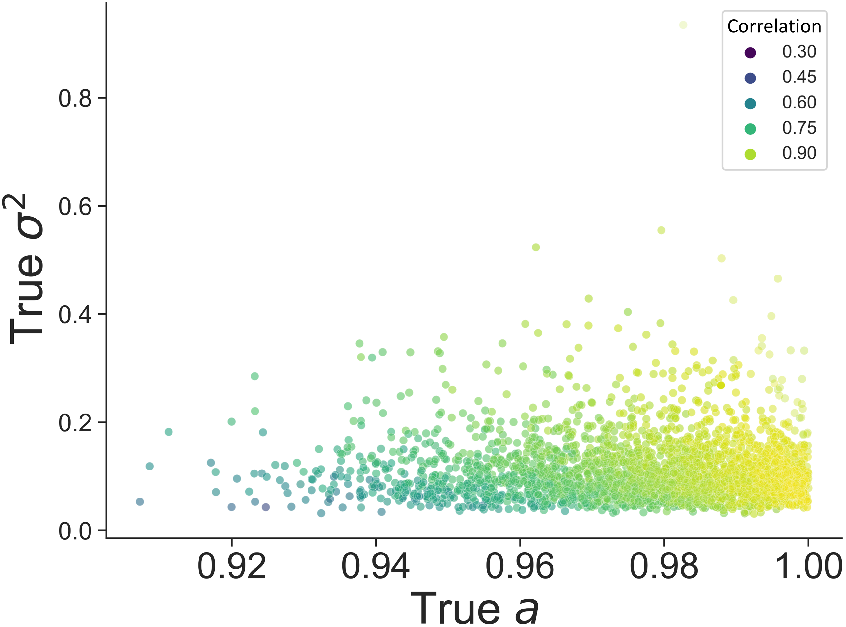
The correlation between the true and the estimated criterion trajectory in function of the true values for *a* and *σ*^2^. The criterion trajectory is harder to recover when lower values for *a* are combined with lower values for *σ*^2^. These parameter values typically generate time series that do not fluctuate much and overall stay relatively close to its baseline, making it harder for the model to identify.

